# Cholesterol and sphingomyelin are critical for Fcγ receptor-mediated phagocytosis of *Cryptococcus neoformans* by macrophages

**DOI:** 10.1101/2021.11.02.466923

**Authors:** Arielle M. Bryan, Jeehyun Karen You, Guangtao Li, JiHyun Kim, Ashutosh Singh, Johannes Morstein, Dirk Trauner, Nívea Pereira de Sá, Tyler G. Normile, Amir M. Farnoud, Erwin London, Maurizio Del Poeta

## Abstract

*Cryptococcus neoformans* is a fungal pathogen that causes life-threatening meningoencephalitis in lymphopenic patients. Pulmonary macrophages comprise the first line of host defense upon inhalation of fungal spores, whereby macrophages either aid in clearance or serve as a niche for its dissemination. Given that macrophages play a key role in the outcome of a cryptococcal infection, it is crucial to understand factors that mediate phagocytosis of *C. neoformans*. Since lipid rafts (high order plasma membrane domains enriched in cholesterol and sphingomyelin) have been implicated in facilitating phagocytosis, we evaluated whether these ordered domains govern macrophages’ ability to phagocytose *C. neoformans*. We found that cholesterol or sphingomyelin depletion resulted in significantly deficient IgG-mediated phagocytosis of the fungus. Moreover, repletion of macrophage cells with a raft-promoting sterol (7-dehydrocholesterol) rescued this phagocytic deficiency while a raft-inhibiting sterol (coprostanol) significantly decreased IgG-mediated phagocytosis of *C. neoformans*. Using a photoswitchable sphingomyelin (AzoSM), we observed that the raft-promoting conformation (*trans*-AzoSM) resulted in efficient phagocytosis whereas raft-inhibiting conformation (*cis*-AzoSM) significantly blunted phagocytosis in a reversible manner. We observed that the effect on phagocytosis may be mediated by facilitating Fcγ receptor (FcγR) function, whereby IgG immune complexes cross-link to FcγRIII, resulting in tyrosine phosphorylation of FcR γ-subunit (FcRγ), an important accessory protein in the FcγR signaling cascade. Correspondingly, cholesterol or sphingomyelin depletion resulted in decreased FcRγ phosphorylation. Repletion with 7-dehydrocholesterol restored phosphorylation, whereas repletion with coprostanol showed FcRγ phosphorylation comparable to unstimulated cells. Together, these data suggest that lipid rafts are critical for facilitating FcγRIII-mediated phagocytosis of *C. neoformans*.

## Introduction

*Cryptococcus* spp. are major opportunistic fungal pathogens that cause life-threatening meningoencephalitis in immunocompromised individuals (1,2). *Cryptococcus neoformans* has been reported to cause approximately 223,100 cases of meningitis globally, resulting in 181,000 deaths every year (3). *C. neoformans* mediated host damage at the molecular and cellular level results in deficient immune effector cell function, disruption of endothelial barriers, and dissemination (4). Despite the substantial morbidity and mortality associated with cryptococcal meningoencephalitis, there is still much that is unknown about the delicate interactions between host immune cells and fungal pathogens like *C. neoformans*.

Pulmonary macrophages comprise the first line of host defense in response to C. *neoformans*, as infection begins with inhalation of fungal spores commonly found in urban environments. A healthy host can typically kill and clear the pathogen. However, the fungus can be a facultative intracellular pathogen that survive within macrophages in immunocompromised individuals (e.g., HIV and AIDS patients) (2,5). In these cases, the macrophages can serve as a niche for the replication of the pathogen and may facilitate its dissemination to the central nervous system where the disease becomes fatal (6–9). *C. neoformans* can impair mitochondrial function, alter protein synthesis, or undergo non-lytic exocytosis from macrophages (10–13). Macrophages may even deliver the fungus directly into the meninges, helping the yeast to cross the blood brain barrier via the “Trojan horse” model (14–17). Given that macrophages can determine the outcome of a cryptococcal infection, it is crucial to understand the factors that mediate phagocytosis of *C. neoformans*, a major function of macrophages.

Phagocytosis is a process by which extracellular entities are internalized by phagocytic cells. It is a key weapon in the immune system’s arsenal to defend against pathogens, but the process may often be subverted by pathogens to allow for internalization and dissemination throughout the body (18). Phagocytosis is mediated by several signaling events that result in attachment and engulfment via rearrangements of the host cell’s cytoskeleton. ‘Professional’ phagocytes are able to recognize and bind to opsonins on the surface of the invading pathogen to signal for attachment and the formation of lamellipodia, which engulf the pathogen and form a phagosome (2). Macrophages are highly specialized cells that carry out protective functions that include seeking out and eliminating disease causing agents, repairing damaged tissues, and mediating inflammation, most of these through the process of phagocytosis (18,19).

Previous work in other pathogen systems point to lipid rafts formed by cholesterol and sphingomyelin as having an important role to play in phagocytosis (20–26). In fact, a recent study implicated lipid rafts in the phagocytic response to *Aspergillus fumigatus*, another opportunistic fungal pathogen (27). Cholesterol is the single most abundant lipid species in mammalian plasma membranes, comprising 25-50% of the cell membrane lipid content (28). It has been found to play a role in modulating the biophysical properties of membranes by changing their rigidity (29). Cholesterol and sphingolipids together form lipid microdomains within the membrane known as lipid rafts. Lipid rafts have been found to be involved in the formation of caveolae as well as activation of various signaling cascades, whereby domains favor specific protein-protein interactions (28–30). In fact, previous studies have implicated the role of rafts in immune receptor signaling (31). Given their small size, it is difficult to study lipid rafts *in vivo*. One useful way to study the role of lipid rafts is to alter their constituents. Cyclodextrins and sphingomyelinase have been found to deplete, replace, or alter lipids from host mammalian membranes and are commonly used to study the role of lipid rafts (30,32–37).

In this study, several approaches were employed to alter cholesterol or sphingomyelin on the outer leaflet of the plasma membrane to examine how lipid rafts may play a role in phagocytosis of *C. neoformans*. We found that cholesterol and sphingomyelin are critical for IgG-mediated phagocytosis of *C. neoformans*. Moreover, repletion with domain-promoting sterols like cholesterol or 7-dehydrocholesterol promoted efficient antibody-mediated phagocytosis of the fungal cells, whereas domain-inhibiting sterols like coprostanol significantly reduced it. Using a photoswitchable sphingomyelin (AzoSM), we observed that the raft-promoting conformation (*trans*-AzoSM) resulted in efficient phagocytosis whereas raft-inhibiting conformation (*cis*-AzoSM) significantly blunted phagocytosis in a reversible manner. Mechanistically, we found that cholesterol and sphingomyelin enriched ordered domains may be important for the function of Fcγ receptors, the class of immune receptors activated by IgG immune complexes. These findings provide more directed insights into the role of cholesterol and sphingomyelin-rich lipid rafts in mediating Fcγ receptor activation and IgG-dependent phagocytosis of *C. neoformans*.

## Results

### Cholesterol or sphingomyelin depletion affects IgG-mediated phagocytosis of C. neoformans

To investigate the role of cholesterol in phagocytosis of the fungal pathogen *C. neoformans*, we utilized methyl-beta-cyclodextrin (MβCD) to deplete cholesterol from the membrane of murine macrophages prior to infection with antibody opsonized *C. neoformans* (38). We previously showed that treatment with 10 mM or 30 mM MβCD depletes approximately 50% or 75% of the total cholesterol in the cells, respectively (38). Furthermore, MβCD treatment did not significantly alter cell attachment or viability (38). Most importantly, cholesterol depletion resulted in a significant decrease in antibody-mediated phagocytosis of *C. neoformans* (38). This finding was recapitulated across two murine macrophage cell lines. Both alveolar (Figure 1A) and peritoneally-derived macrophages (J774.1, Figure S1A) showed significant deficiency in phagocytosing *C. neoformans* cells opsonized with an anti-glucuronoxylomannan (GXM) IgG, an antibody specific to the cryptococcal capsule (39). When *C. neoformans* cells were instead opsonized with complement serum, phagocytosis was not affected (Figure 1C),

**Figure 1.**
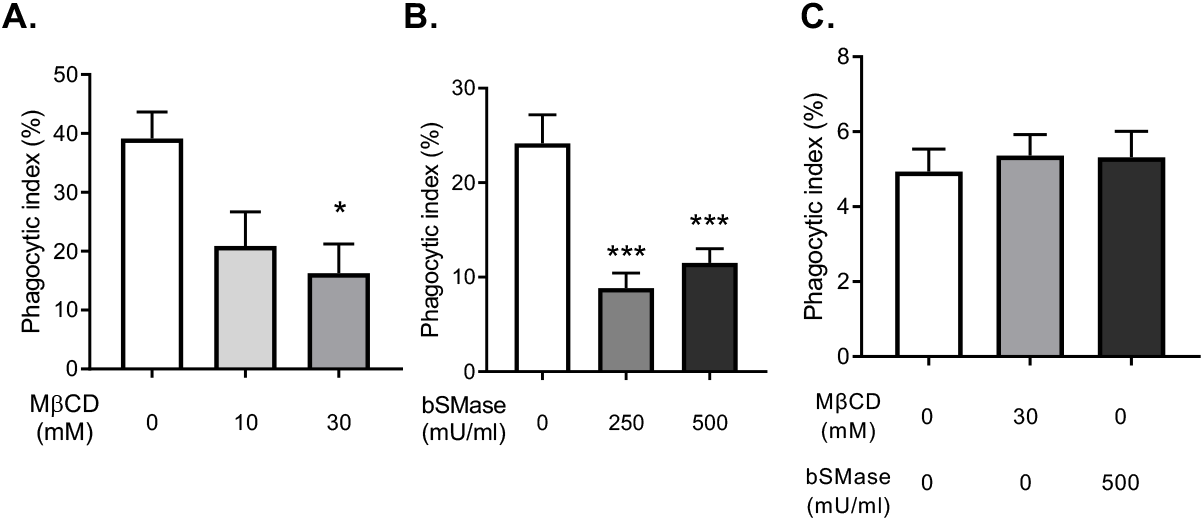
Cholesterol and sphingomyelin are important for antibody-mediated phagocytosis. **A)** Macrophages (MH-S) treated with either 10 or 30 mM methyl-beta-cyclodextrin (MβCD) were co-incubated with antibody-opsonized *C. neoformans* H99 at a 1:1 ratio and allowed to interact for 2 hours. Cells were then fixed and stained with Giemsa and phagocytic index was calculated by microscopic observation (n=4). **B)** Macrophages (MH-S) treated with either 250 or 500 mU/ml bacterial sphingomyelinase (bSMase) were coincubated with antibody-opsonized *C. neoformans* H99 at a 1:1 ratio and allowed to interact for 2 hours. Cells were then fixed and stained with Giemsa and phagocytic index was calculated by microscopic observation (n=3).**C)** Macrophages (MH-S) treated with 30 mM MβCD or 500 mU/ml bSMase were co-incubated with complement serum-opsonized *C. neoformans* H99 at a 1:1 ratio and allowed to interact for 3 hours. Cells were then fixed and stained with Giemsa and phagocytic index was calculated by microscopic observation (n=4). Error bars represent the standard error of the mean (SEM), and statistical significance was determined using one-way ANOVA with Tukey’s multiple comparisons test. \**P* < 0.05, \*\*\**P* < 0.001 compared to the untreated control. All *P*-values were adjusted for multiplicity.

Given the collaborative role of cholesterol and sphingomyelin in lipid rafts, we examined the effect of depleting sphingomyelin on the plasma membrane of macrophages. One tool available for the study of sphingomyelin is recombinant bacterial sphingomyelinase (bSMase) which directly probes for the role of sphingomyelin on the plasma membrane, as the enzyme is too large to pass through the membrane (40). bSMase catalyzes the transformation of sphingomyelin into ceramide (Figure S2) and phosphorylcholine (40). To confirm sphingomyelin depletion, cellular lipids were analyzed following treatment with 250 mU/ml or 500 mU/ml bSMase for 20 minutes. We found that treatment of macrophages with bSMase resulted in a significant decrease in C16 sphingomyelin, the most abundant sphingomyelin species detected in the cells, and a corresponding increase in C16 ceramide (Figure 2). To assess the effect of bSMase treatment on phagocytosis, cells were co-incubated with *C. neoformans* cells opsonized with anti-GXM IgG after bSMase treatment. We found a significant decrease in phagocytosis after bSMase treatment with both alveolar (MH-S, Figure 1B) and peritoneally-derived macrophages (J774.1, Figure S1B), albeit in a non-dose-dependent fashion. This effect was not observed when complement serum was used as an opsonin (Figure 1C). Together these data suggest that cholesterol and sphingomyelin are critical for IgG-mediated phagocytosis of *C. neoformans*. Moreover, our data suggest that this phenomenon is not cell line dependent.

**Figure 2.**
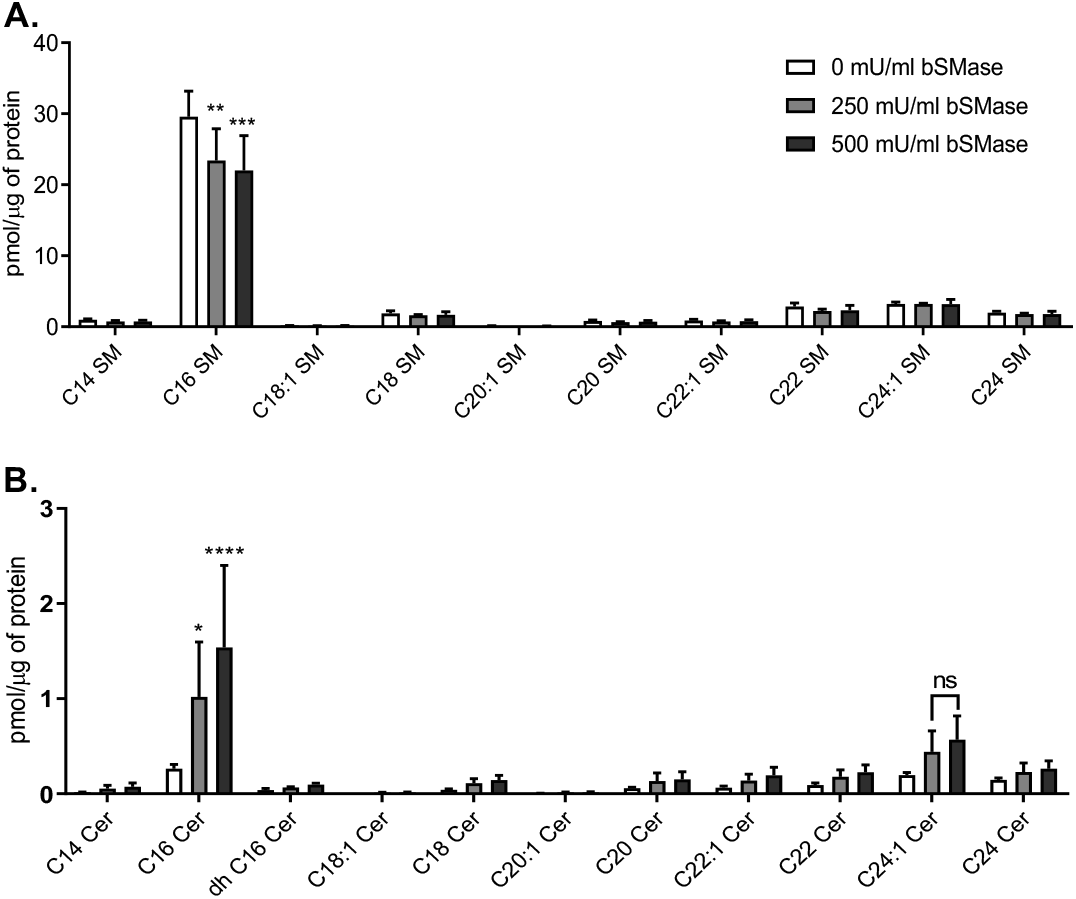
Measurement of sphingolipids after treatment with bacterial sphingomyelinase. Macrophages (J774.1) were treated with either 250 or 500 mU/ml bacterial sphingomyelinase (bSMase). Monolayers were then washed and subject to lipid extraction. Lipid extracts were then analyzed using liquid chromatography coupled mass spectrometry and normalized to protein found in the cell extract using the Bradford assay. **A)** Sphingomyelin (SM) species in the treated samples are shown (n=3). **B)** Ceramide (Cer) species in the treated samples are shown (n=3). Error bars represent the SEM and statistical significance was determined using ANOVA with Tukey’s multiple comparisons test. Not significant (ns), \**P* < 0.05, \*\**P* < 0.01, \*\*\**P* < 0.005, \*\*\*\**P* < 0.0001 compared to the untreated control. All *P*-values were adjusted for multiplicity.

### Repletion with lipid raft altering sterols affects IgG-mediated phagocytosis of C. neoformans

Given that multiple factors confound the use of MβCD to deplete cholesterol from cell membranes, various sterols were added back into the cholesterol depleted macrophages to investigate the role of cholesterol in IgG-mediated phagocytosis of *C. neoformans*. MβCD may remove cholesterol from both raft and non-raft domains, alter the distribution of cholesterol between plasma and intracellular membranes, and non-specifically extract phospholipids (41). To ascertain whether cholesterol sensitivity of IgG-dependent phagocytosis of *C. neoformans* could be attributed to lipid rafts, cholesterol depleted macrophages were repleted with cholesterol, 7-dehydrocholesterol, or coprostanol (Figure S2). 7-dehydrocholesterol has been shown to be significantly more domain-promoting than cholesterol, whereas coprostanol strongly inhibits domain formation (36,42,43). We found that repletion with 0.2 mM cholesterol in 2.5 mM MβCD resulted in a significant increase in total cellular cholesterol compared to the untreated control. On the other hand, repletion with 0.2 mM 7-dehydrocholesterol or 0.2 mM coprostanol in 2.5 mM MβCD resulted in a significant decrease in cellular cholesterol and a marked increase in substituted sterol comparable to the cellular cholesterol for the untreated control (Figure 3A). When treated macrophages were co-incubated with *C. neoformans* cells opsonized with anti-GXM IgG, we found that we were able to restore phagocytosis by cholesterol or 7-dehydrocholesterol repletion, whereas coprostanol repletion did not restore phagocytosis (Figure 3B).

**Figure 3.**
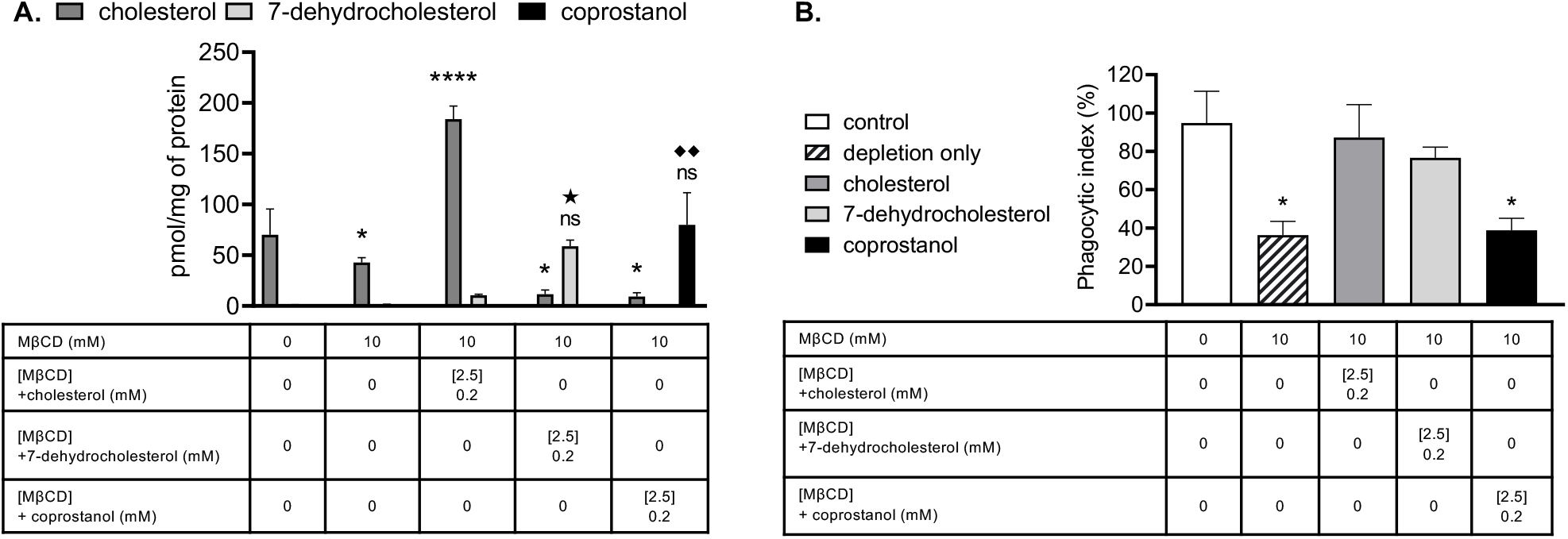
Repletion with raft altering sterols affects phagocytosis. **A)** Macrophages (J774.1) were pre-treated with 10 mM MβCD to deplete cholesterol and then washed and incubated with 2.5 mM MβCD loaded with 0.2 mM of indicated sterol. Monolayers were then washed and subject to lipid extraction. Lipid extracts were then analyzed using gas chromatography coupled mass spectrometry and normalized to protein found in the cell extract by the Bradford assay (n=3). **B)** Macrophages were pre-treated with 10 mM MβCD to deplete cholesterol and then washed and incubated with 2.5 mM MβCD loaded with 0.2 mM of indicated sterol. Monolayers were then washed and allowed to interact with antibody opsonized *C. neoformans* H99 at a 1:1 ratio for 2 hours. Cells were then fixed and stained with Giemsa and phagocytic index was calculated by microscopic observation (n=3). Error bars represent SEM and statistical significance was determined using one-way ANOVA with Tukey’s multiple comparisons test. Not significant (ns), \**P* < 0.05, \*\**P* < 0.01, \*\*\*\**P* < 0.0001 compared to the cellular cholesterol for untreated control. ^★^*P* < 0.05 compared to the cellular 7-dehydrocholesterol for untreated control. ^uu^*P* < 0.01 compared to the cellular coprostanol for untreated control. All *P*-values were adjusted for multiplicity.

### Cholesterol, but not sphingomyelin digestion affects lipid nanodomain stability

To assess how sterol depletion/repletion may affect lipid raft stability in macrophages, we evaluated ordered nanodomains (rafts) in giant plasma membrane vesicles (GPMVs) derived from macrophages through Förster resonance energy transfer (FRET). When rafts are present FRET donor and FRET acceptor become partially segregated in different (raft and non-raft) domains, and FRET decreases so the FRET donor is more fluorescent (i.e. F/F_0_ increases) As previously, we used the temperature at the which the value of F/F_0_ is a minimum as an approximate temperature for the upper limit of when rafts were present, i.e. a measure of their thermal stability (36). We also used the total increase in F/F_0_ relative to the value at which F/F_0_ is a minimum in a sample as another rough measure of total raft formation over the entire temperature range.

GPMVs prepared from cholesterol-depleted macrophages showed a significant shift in nanodomain stability (Figure 4A, Table S1, raw unnormalized F/F_0_ values for FRET data are shown in Figures S3A). The presence of detectable ordered nanodomains significantly decreased, with up to 20 °C decrease in T_end_, the temperature at which ordered domains are completely melted (36). Upon repletion with cholesterol, detectable ordered nanodomains significantly increased compared to the untreated and cholesterol depleted macrophages with a corresponding recovery in T_end_. As expected, repletion with 7-dehydrocholesterol resulted in significantly greater presence of detectable nanodomains as well as increased thermal stability (higher T_end_ compared to untreated control). On the other hand, repletion with coprostanol ablated both the presence of detectable ordered nanodomains and the GPMV thermal stability (Figure 4B, Table S1, raw unnormalized F/F_0_ values for FRET data are shown in Figure S3B).

**Figure 4.**
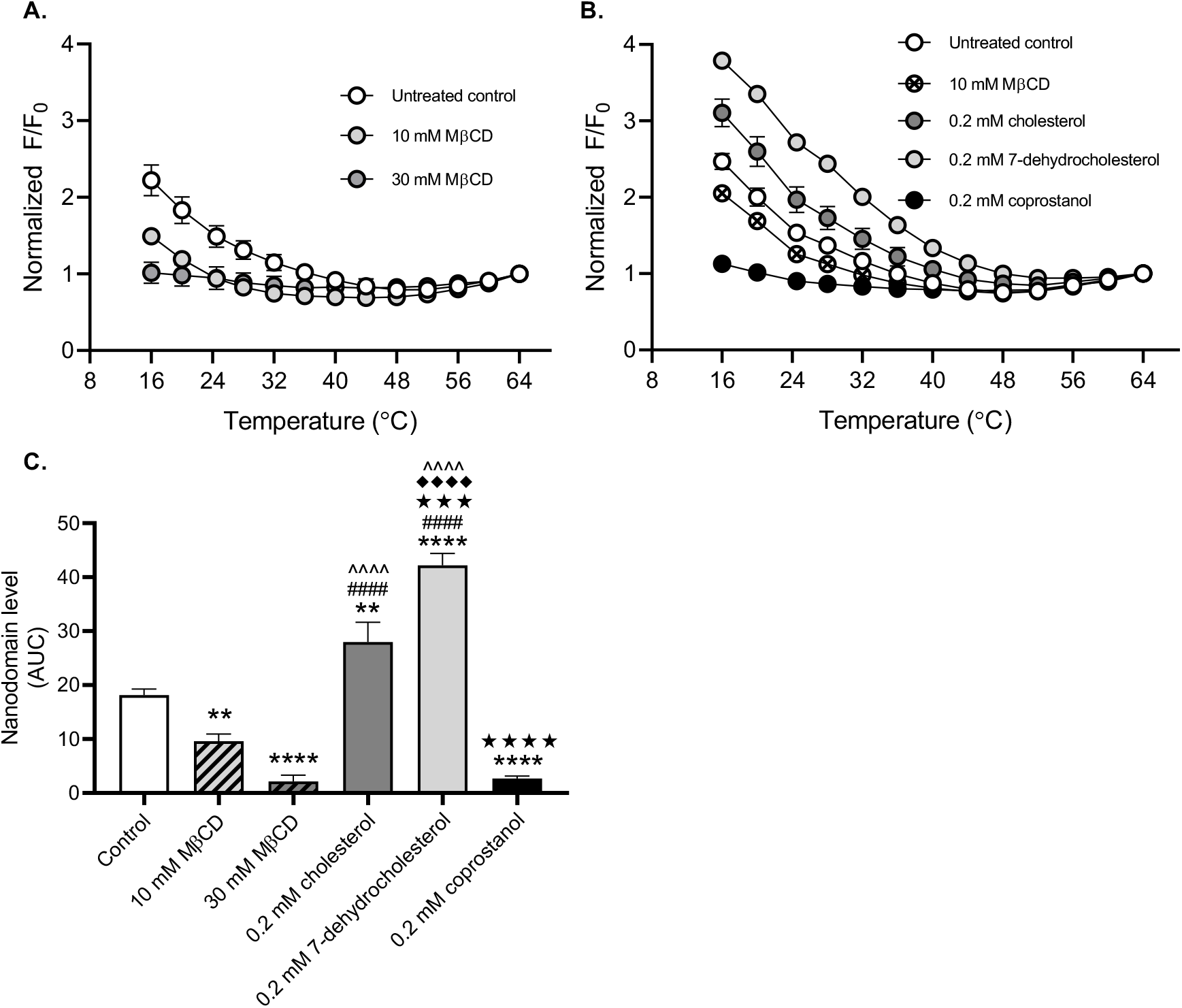
Repletion with raft altering sterols affects nanodomain stability. **A)** Macrophages (MH-S) were pre-treated with either 10 or 30 mM MβCD to deplete cholesterol. Monolayers were then washed and subject to giant plasma membrane vesicle formation using 25 mM PFA and 2 mM DTT. Nanodomain stability was assessed via FRET with DPH as the FRET donor and ODRB as the FRET acceptor. The ratio of DPH fluorescence intensity in the presence vs. absence of ODRB was calculated (F/F_0_). F/F_0_ values were normalized to the final F/F_0_ value at 64°C (n=3). **B)** Macrophages were pre-treated with 10 mM MβCD to deplete cholesterol and then washed and incubated with 2.5 mM MβCD loaded with 0.2 mM of indicated sterol. Monolayers were then washed and subject to giant plasma membrane vesicle formation using 25 mM PFA and 2 mM DTT. Nanodomain stability was assessed via FRET with DPH as the FRET donor and ODRB as the FRET acceptor. The ratio of DPH fluorescence intensity in the presence vs. absence of ODRB was calculated (F/F_0_). F/F_0_ values were normalized to the final F/F_0_ value at 64°C (n=3). **C)** Relative domain levels were estimated using the polynomial fits from A & B as described under *Experimental procedures*. Error bars represent SEM. Detectable nanodomains were compared using one-way ANOVA with Tukey’s multiple comparisons test. \*\**P* < 0.01, \*\*\*\**P* < 0.0001 compared to the untreated control, ^####^*P* < 0.0001 compared to 10 mM MβCD, ^^^^*P* < 0.0001, ^★★★^*P* < 0.001, ^★★★★^*P* < 0.0001 compared to 0.2 mM cholesterol, and ^◆◆◆◆^*P* < 0.0001 compared to 0.2 mM coprostanol. All *P*-values were adjusted for multiplicity.

To ascertain whether sphingomyelin depletion alters lipid raft stability in macrophages, we also evaluated nanodomain stability in GPMVs derived from bSMase-treated macrophages. We found that neither 250 nor 500 mU/ml bSMase treatment altered nanodomain stability, with T_end_ and GPMV levels comparable to the untreated control (Figure S4, Table S1, raw unnormalized F/F_0_ values for FRET data are shown in Figure S4C). It should be noted that the lipid product of bSMase digestion, ceramide, is itself a raft-forming lipid, but one that alters the lipid composition (e.g. resulting in displacement of cholesterol from rafts) and properties of lipid rafts (44,45).

Together these results not only suggest that the presence of ordered lipid domains mediate IgG-dependent phagocytosis of *C. neoformans*, but that lipid domain stability is cholesterol-dependent.

### Cholesterol or sphingomyelin depletion may affect structure of lipid rafts

EGFP-nakanori is a protein that labels cell surface domains in a sphingomyelin and cholesterol-dependent manner. More specifically, nakanori only binds pre-existing sphingomyelin/cholesterol domains as it is unable to induce formation of sphingomyelin-cholesterol complexes (46). To examine whether cholesterol depletion or sphingomyelin depletion perturbs sphingomyelin-cholesterol complexes, macrophages were either treated with MβCD or bSMase and labeled with EGFP-nakanori. Both cholesterol and sphingomyelin depletion resulted in marked decrease in EGFP-nakanori binding, suggesting successful perturbation of sphingomyelin-cholesterol complexes (Figure 5). These results suggest that MβCD and bSMase treatment may potentially affect lipid raft structure independent of nanodomain stability.

**Figure 5.**
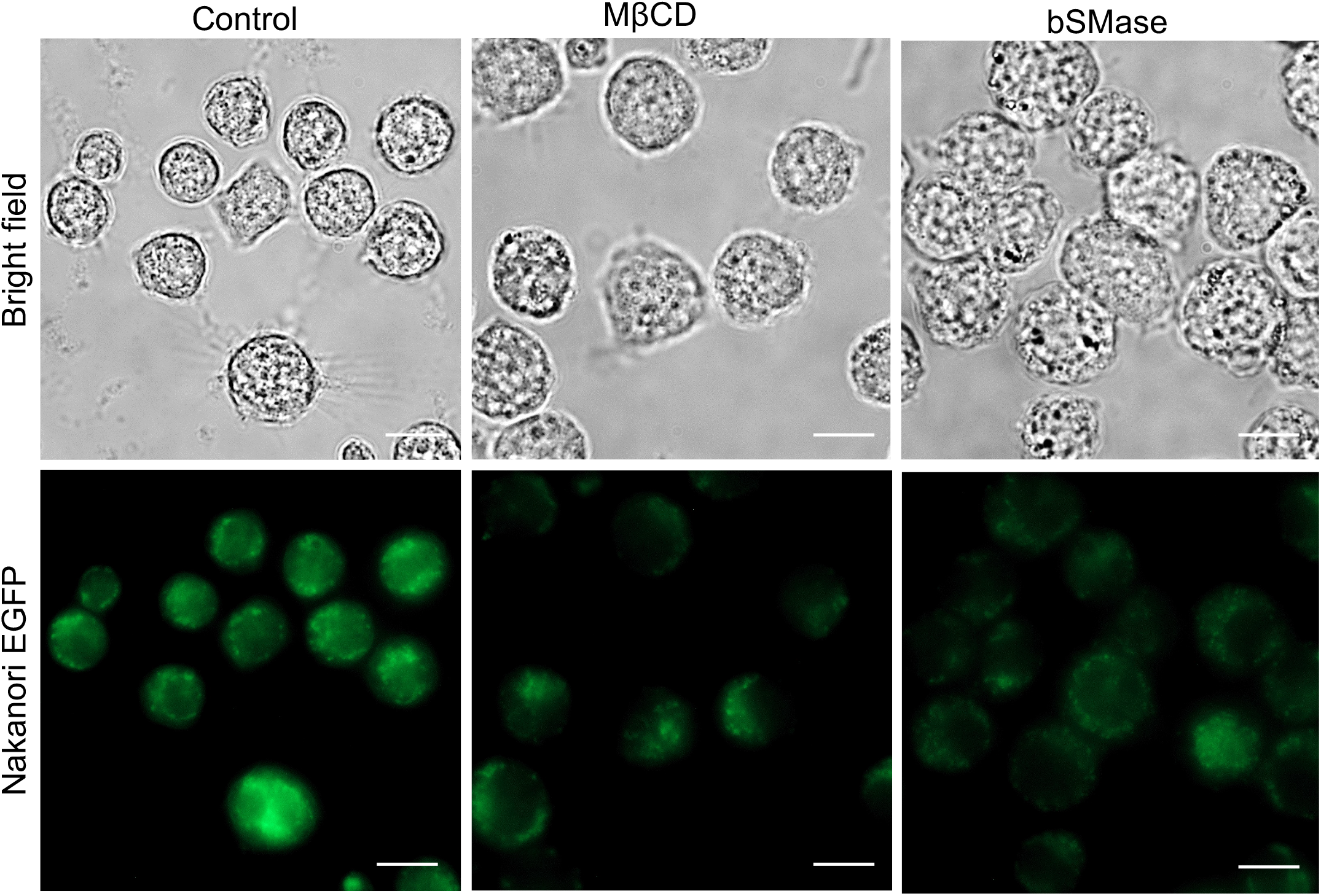
Nakanori labeling is affected by lipid altering treatments. Macrophages (MH-S) were adhered onto glass bottomed microscopy dishes. After treatment with 10 mM MβCD or 250 mU/ml bacterial sphingomyelinase (bSMase) as described under *Experimental Procedures*, live macrophages were labeled with recombinant nakanori-EGFP and then fixed with 4% paraformaldehyde. Scale bar=10 μm.

### AzoSM can reversibly modulate IgG-mediated phagocytosis of C. neoformans

To better understand how sphingomyelin facilitates IgG-dependent phagocytosis of *C. neoformans*, we utilized a photoswitchable sphingomyelin. Photoswitchable lipids allow for the optical control of lipid function with high spatiotemporal resolution, enabling precise functional manipulations. They have been shown to modulate many aspects of membrane biophysics, including permeability, fluidity, lipid mobility and domain formation (47–49). Recently, photoswitchable sphingolipids have emerged as potent tools to investigate cellular physiology using optical control, including receptor-mediated signaling and immune function (50–53). We synthesized AzoSM, a C16:0 sphingomyelin derivative containing an isosteric azobenzene photoswitch following a design principle called ‘azologization’. Optical control of photoswitchable sphingolipids is mediated through light exposure. UV light (λ=365 nm) isomerizes the azobenzene group from a *trans* to *cis* conformation whereas blue light (λ=460 nm) reverts ~80% of AzoSM from a *cis* to *trans* conformation (Figure S2). UV light therefore introduces a kink into the fatty acyl tail to disrupt ordered domains while blue light promotes domain formation (48,56). Using methyl-alpha-cyclodextrin (MαCD), endogenous sphingomyelin species in macrophages were exchanged for AzoSM. This method exchanges phospholipids but does not alter cell sterol (32). We found that MαCD effectively replaced ~80% of endogenous C16:0 sphingomyelin, the most abundant species of sphingomyelin in MH-S murine macrophages, with AzoSM (Figure 6A). When macrophages were coincubated with *C. neoformans* cells opsonized with anti-GXM IgG immediately following AzoSM exchange (no light exposure), we observed phagocytosis comparable to untreated control cells. Interestingly, we found that UV light treatment prior to co-incubation with *C. neoformans* resulted in a significant decrease in phagocytosis. Moreover, UV light exposure immediately followed by blue light treatment prior to co-incubation with fungal cells partially restored phagocytosis (Figure 6B). Together these data suggest that sphingomyelin-dependent lipid domains may be critical for IgG-mediated phagocytosis of *C. neoformans*.

**Figure 6.**
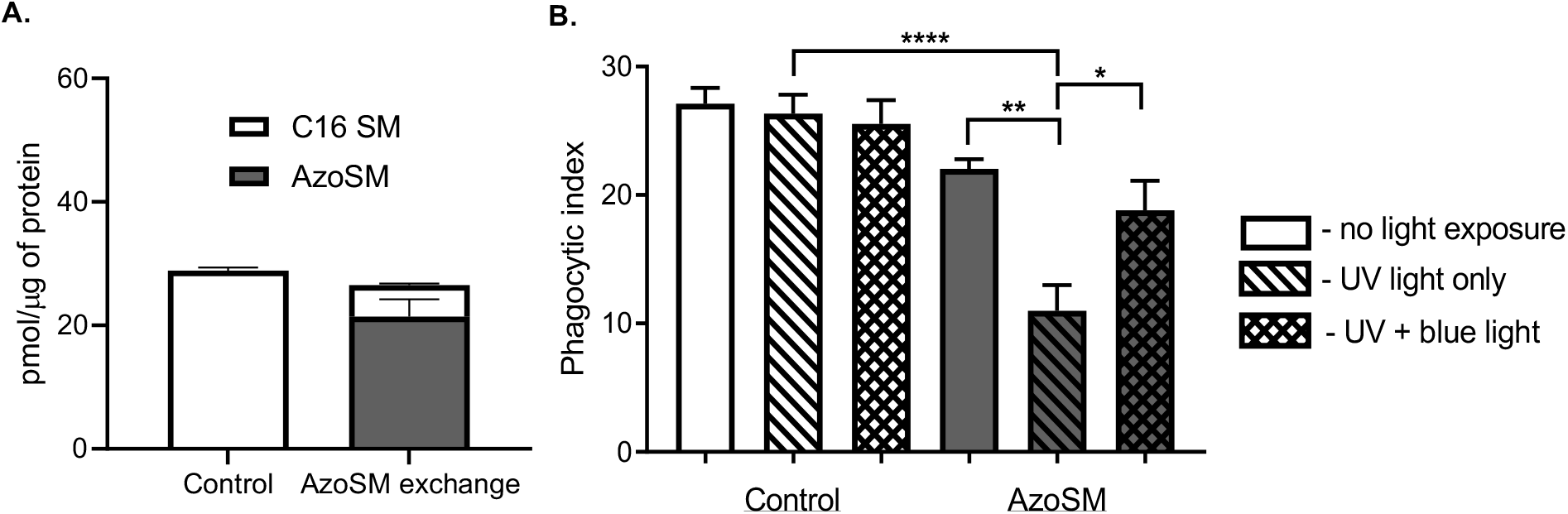
Exclusion of sphingomyelin from rafts affects phagocytosis *C. neoformans*. **A)** Macrophages (MH-S) treated with the AzoSM exchange protocol using MαCD as described in *Experimental Procedures*. Cell extracts were then evaluated for endogenous C16 sphingomyelin (C16 SM) using LC-MS and AzoSM using HPLC. Lipid content from each sample was normalized by protein content as determined using the Bradford assay (n=3) **B)** Macrophages (MH-S) treated with the AzoSM exchange protocol were either unexposed, exposed to UV light, or exposed to UV light followed by blue light. Control samples were mock treated. Washed monolayers were then co-incubated at a 1:1 ratio with *C. neoformans* H99 opsonized with 18B7 antibody and allowed to interact for 30 minutes. Cells were then fixed and stained with Giemsa and phagocytic index was calculated by microscopic observation. Error bars represent the standard error of the mean and statistical significance was determined using two-way analysis of variance with Tukey’s multiple comparisons test (n=4). \**P* < 0.05, \*\**P* < 0.01, and \*\*\*\**P* < 0.0001. All *P*-values were adjusted for multiplicity.

**Figure 7.**
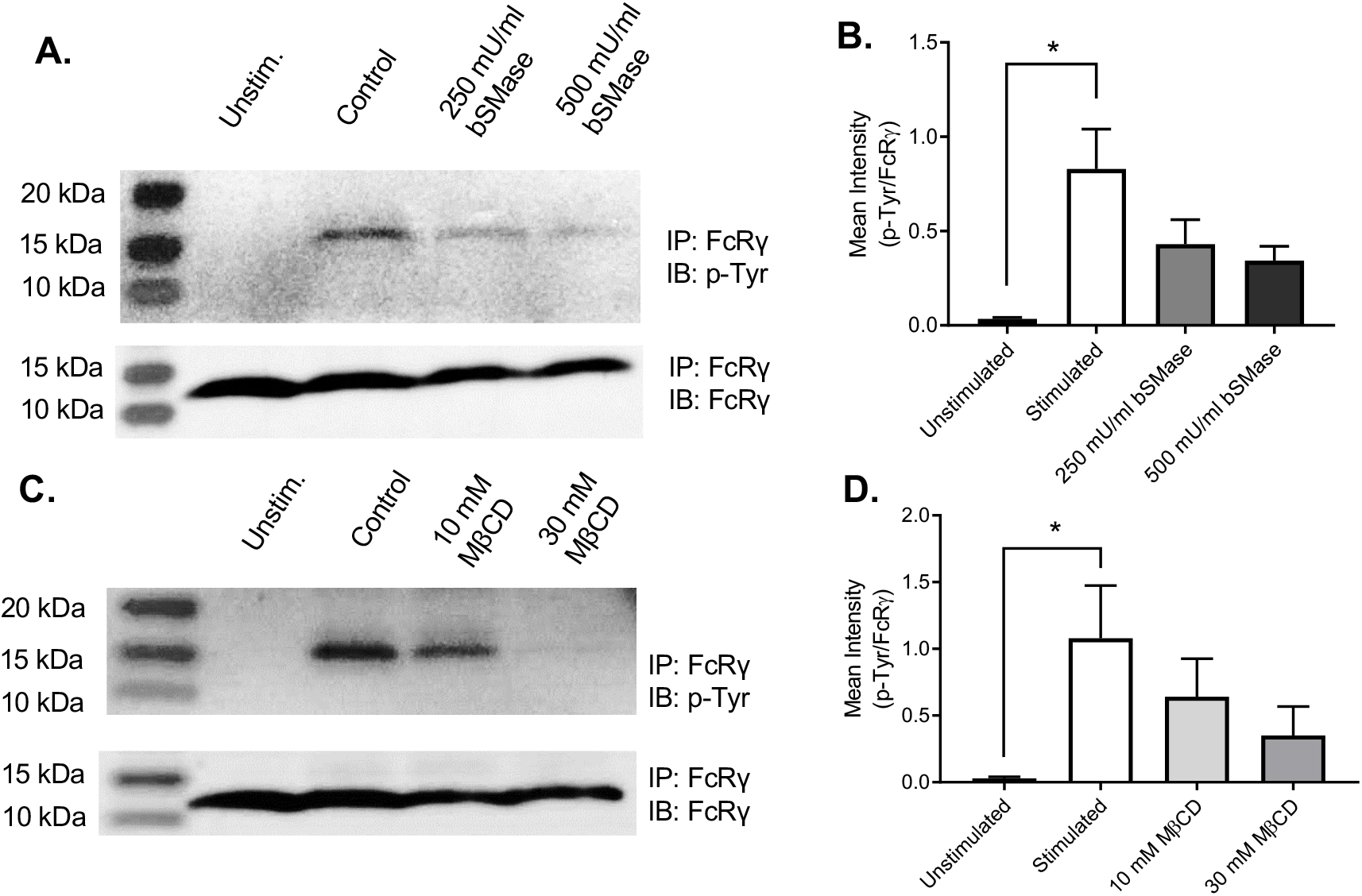
Lipid depleting treatments affect Fcγ receptor-mediated signaling. **A)** Macrophages (MH-S) were mock-treated (control) or treated with bSMase and subsequently stimulated with BSA-IgG immune complex (IC) for 5 min. Fc receptor γ chain (FcRγ) was immunoprecipitated (IP) and evaluated for phosphorylation of the tyrosine residues. IP samples were probed for FcRγ or phospho-tyrosine (p-Tyr). Representative immunoblots are shown. **B)** Immunoblots were quantified using ImageJ software; p-Tyr was normalized by FcRγ (n=3 experiments). Error bars represent the standard error of the mean (SEM), and statistical significance was determined using one-way ANOVA with Tukey’s multiple comparisons test. \**P* < 0.05. All *P*-values were adjusted for multiplicity using Bonferroni’s correction. **C)** MH-S cells were mock-treated (control) or treated with MβCD and subsequently stimulated with BSA-IgG IC for 5 min. FcRγ was IP and evaluated for phosphorylation of the tyrosine residue. IP samples were probed for FcRγ or p-Tyr. Representative immunoblots are shown. **D)** Immunoblots were quantified using ImageJ software; p-Tyr was normalized by FcRγ (n=3). Error bars represent the SEM and statistical significance was determined using one-way analysis of variance with Tukey’s multiple comparisons test compared to the control. \**P* < 0.05. All *P*-values were adjusted for multiplicity.

### Alterations in rafts affect Fcγ receptor function

Since alterations in plasma membrane lipids only affected phagocytosis of *C. neoformans* opsonized with anti-GXM IgG, our data implicate a critical role for lipid rafts in Fcγ receptor (FcγR) mediated phagocytosis. As the anti-GXM antibody falls in the IgG1κ subclass, antibody-opsonized *C. neoformans* bind to FcγRIIB and III, the low-affinity receptors for moue IgG1 (39,57). To assess whether lipid depletion treatments alter the amount of cell surface receptors, macrophages were probed for CD16/32, which recognizes FcγR II/III, and CD11b antibody, which recognizes complement receptor 3, known to mediate uptake of serum opsonized *C. neoformans* (58). CD45 was used as a general leukocyte marker, which should not be affected by lipid altering treatments. Flow cytometry showed that neither CD16/32 or CD11b levels on the cell surface were affected by cholesterol depletion with MβCD or sphingomyelin depletion with bSMase for either J774.1 or MH-S macrophages (Figure S5). We next probed whether FcγR function rather than abundance on the plasma membrane was affected by lipid depletion. Murine FcγRIII requires the association of the FcR γ subunit (FcRγ) following IgG immune complex and receptor aggregation at the plasma membrane. The FcRγ contains an immunoreceptor tyrosine-based activation motif (ITAM) that is required for cell activation (57). To investigate FcRγ activation following IgG immune complex and FcγR cross-linking, we utilized the approach by Rittirsch et al (59). We observed tyrosine phosphorylation of FcRγ in macrophages stimulated with BSA-IgG immune complexes (IgGICs) within 5 minutes, with the phospho-tyrosine levels subsiding 15 minutes after stimulation (Figure S6). We then probed for tyrosine phosphorylation of FcRγ in macrophages following lipid altering treatments. We observed that both sphingomyelin depletion (Figure 5A & B) and cholesterol depletion (Figure 5C & D) resulted in reduced FcRγ activation upon IgGIC stimulation. Interestingly, repletion with cholesterol or 7-dehydrocholesterol showed comparable FcRγ activation to the untreated control, whereas repletion with coprostanol significantly ablated the FcRγ activation upon IgGIC stimulation (Figure 6). Although nuanced, we also observed some decrease in FcR γ chain phosphorylation following UV light exposure post-AzoSM exchange, which was partially rescued upon blue light exposure (Figure S7).

Together, these data suggest that cholesterol and sphingomyelin-dependent lipid rafts are critical for proper FcγR clustering upon IgG immune complex cross linking in order to initiate the FcRγ-mediated signaling cascade essential for phagocytosis of IgG1-opsonized *C. neoformans* (60).

## Discussion

In this study, we utilized various methods of lipid alteration to probe for the role of lipid rafts in phagocytosis of *C. neoformans*. Substituting either endogenous cholesterol with a raft inhibiting sterol (coprostanol) or endogenous sphingomyelin with domain perturbing sphingomyelin (*cis*-AzoSM) lead to a decrease in antibody-mediated phagocytosis of *C. neoformans* (Figure 8). Since depletion treatments perturbed sphingomyelin-cholesterol complexes and significantly affected the FcγR-mediated cell signaling (as evidenced by decreased FcRγ ITAM activation) without altering the amount of FcγR IIB and III localized to the cell surface, it appears that sphingomyelin and cholesterol-dependent lipid rafts are critical for facilitating FcγR-mediated cell signaling and function. Lipid rafts do not appear to be critical for phagocytosis mediated by complement serum opsonization, which was not affected by cholesterol depletion or bSMase treatment. The importance of cholesterol-dependent lipid rafts in FcγR function was further highlighted by the fact that repletion with cholesterol or 7-dehydrocholesterol (raft promoting sterols) resulted in proper FcRγ ITAM activation, whereas repletion with coprostanol (raft inhibiting sterol) significantly reduced FcRγ ITAM activation. Similarly, Indeed, a recent review highlights the importance of plasma membrane domains on IgG Fc receptor function (61). However, studies of FcγRIII have been limited to the human FcγRIIIa expressed on natural killer cells. Furthermore, studies have been limited to cellular fractionation assays and confocal microscopy (62,63). Although murine and human FcγR differ in terms of binding abilities and expression pattern, the function and signaling mechanism are conserved (57). As such, insights into how cholesterol and sphingomyelin, two key components of lipid rafts, alter membrane properties and affect FcγR function can have broader implications.

**Figure 8.**
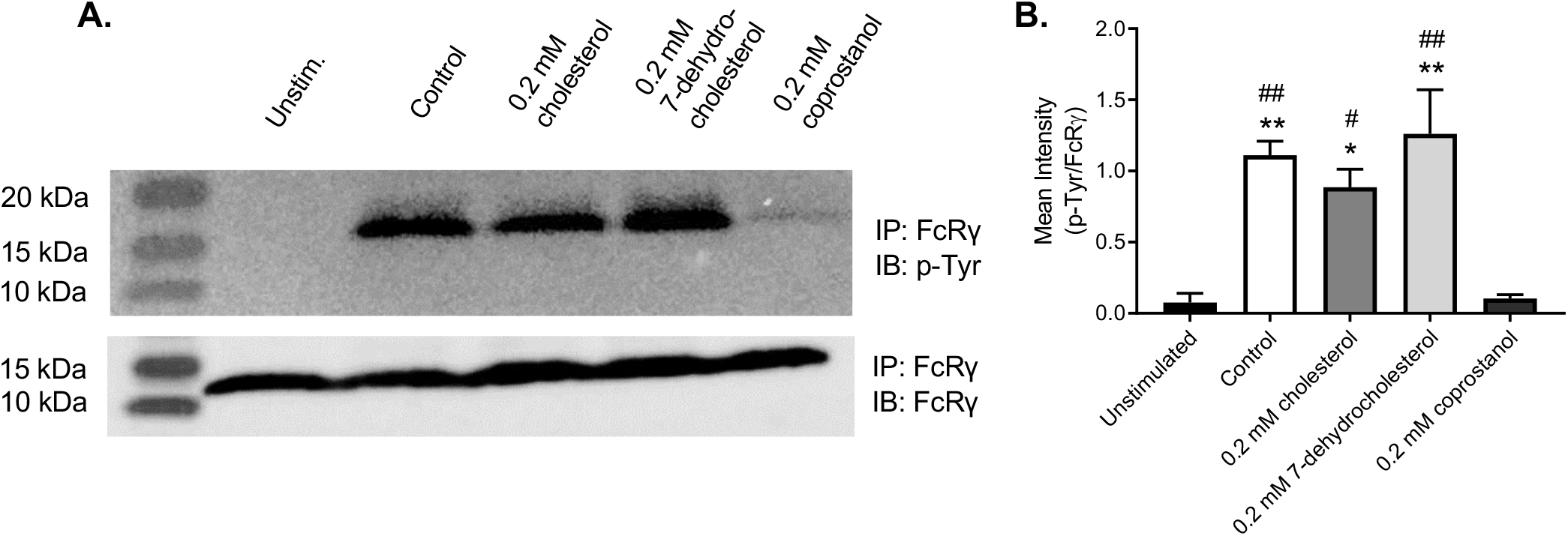
Repletion with raft-altering sterols affects Fcγ receptor-mediated signaling. **A)** Macrophages were mock-treated (control) or pre-treated with 10 mM MβCD to deplete cholesterol and then washed and incubated with 2.5 mM MβCD loaded with 0.2 mM of indicated sterol. Cells were then stimulated with BSA-IgG immune complex (IC) for 5 min. Fc receptor γ chain (FcRγ) was immunoprecipitated and evaluated for phosphorylation of the tyrosine residue. IP samples were probed for FcRγ or phospho-tyrosine (p-Tyr). Representative immunoblots are shown. **B)** Immunoblots were quantified using ImageJ software; p-Tyr was normalized by FcRγ (n=3). Error bars represent the standard error of the mean and statistical significance was determined using one-way ANOVA with Tukey’s multiple comparisons test. \**P* < 0.05, \*\**P* < 0.01, compared to the untreated control. ^#^*P* < 0.05, ^##^*P* < 0.01 compared to 0.2 mM coprostanol. All *P*-values were adjusted for multiplicity.

**Figure 9.**
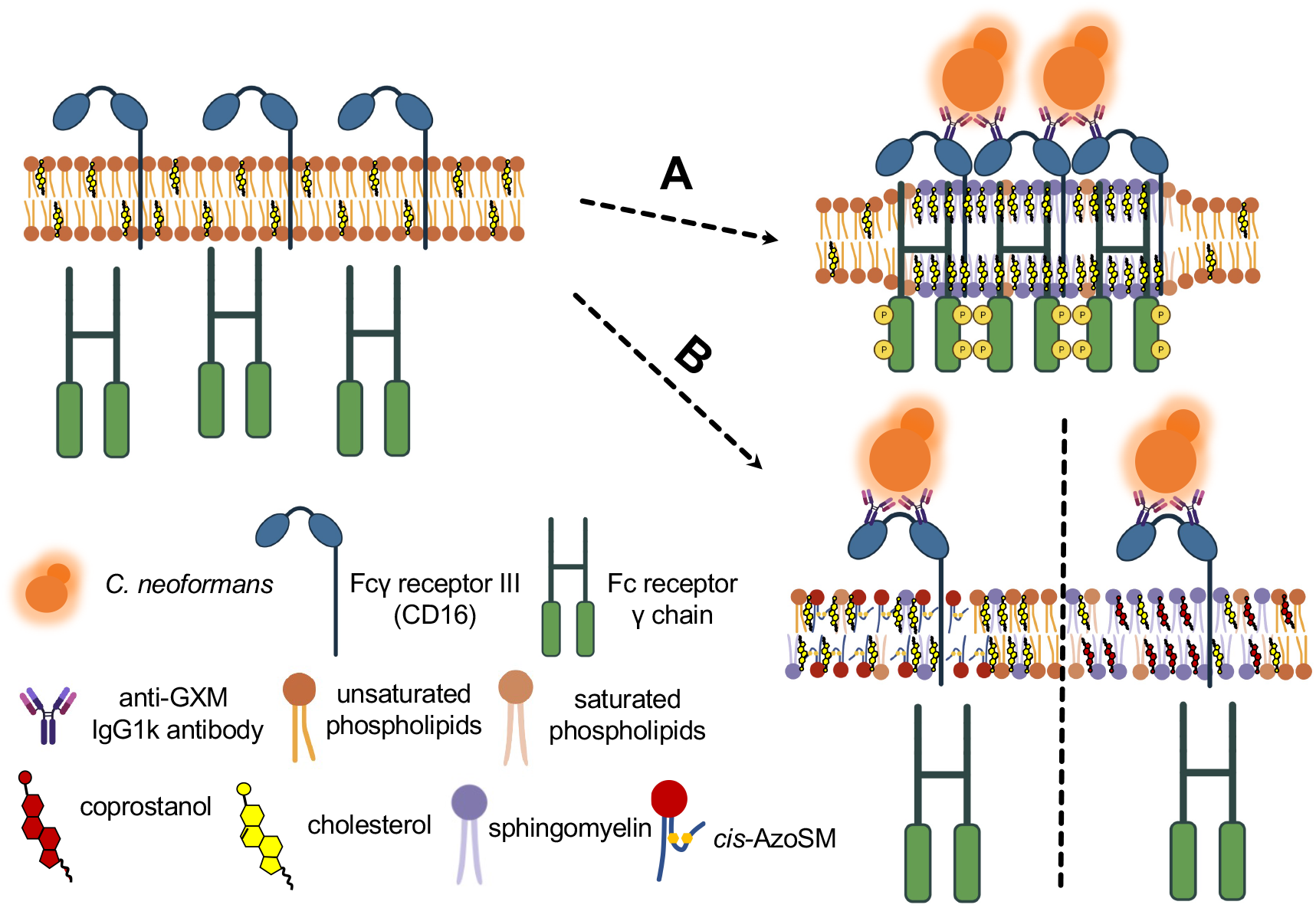
Cholesterol and sphingomyelin rich plasma lipid rafts are important for Fcγ receptor mediated phagocytosis of *C. neoformans* by macrophages. Upon binding IgG-based immune complexes comprised of anti-GXM antibody opsonized *C. neoformans*, Fcγ receptor III (CD16) localizes to the plasma lipid rafts. **A)** When lipid rafts are enriched with raft-forming sterols (i.e., cholesterol and 7-dehydrocholesterol; only cholesterol shown for simplicity) or sphingomyelin (i.e., *trans*-AzoSM; only endogenous sphingomyelin shown for simplicity), the Fc receptor γ chain is properly phosphorylated to initiate the signaling cascade associated with phagocytosis. **B)** When lipid rafts are enriched with raft-inhibiting sterols such as coprostanol (right) or raft-disrupting sphingomyelin, *cis*-AzoSM (left), Fc receptor γ chain is not properly phosphorylated.

Lipid depletion is useful to investigate the role of key lipids in cellular processes. Given the physiological importance of cholesterol in the cell plasma membrane, MβCD has become the most common means of modifying the cholesterol content, mainly through cholesterol extraction. However, cyclodextrins may remove cholesterol from both raft and non-raft domains, alter the distribution of cholesterol between plasma and intracellular membranes, and simultaneously extract phospholipids (41). As such, cholesterol substitution with sterol structural analogues has been utilized in several studies to evaluate specific sterol-protein interactions or changes to the physical properties of the lipid bilayer (33,42,43,64,65). The key is substitution with an equivalent amount of another sterol without altering the total sterol level in cells (41). Although our cholesterol repletion increased total cellular sterol, it did not interfere with macrophages’ ability to phagocytose or initiate FcγR signaling. When compared to 7-dehyrocholesterol or coprostanol repletion, nanodomain properties were consistent with raft formation.

bSMase is a useful tool to study the role of sphingomyelin depletion as well as ceramide accumulation on the plasma membrane. The effects of sphingomyelin digestion can be complex. Ceramides have been implicated in stabilizing sphingomyelin-rich domains (43). They can also displace cholesterol from rafts and from the plasma membrane (44,68,69). Modifying ordered domain properties, therefore, can have pleiotropic consequences. Previous work in other pathogen systems suggested that ceramide-rich membrane rafts resulting from bSMase activity are critical for host defense, including receptor clustering (70,71). Interestingly, ceramide-rich membrane rafts have also been implicated in FcγR signaling. Indeed, the presence of C16 ceramide at the cell surface has been suggested to precede and control FcγRII clustering and phosphorylation in rafts, acting as a negative regulator of IgG-dependent phagocytosis (72,73). FcγRII is an inhibitory immune receptor containing an immunoreceptor tyrosine-based inhibitory motif (ITIM) (74). Given our bSMase treatment resulted in C16 SM depletion and a corresponding accumulation of C16 ceramide (Figure 2), aberrant FcγRII signaling may have contributed to the decrease in IgG-dependent phagocytosis of *C. neoformans*.

Given the complex, pleiotropic effects of bSMase treatment, AzoSM is a potent tool that permits direct control of lipid function and ordered domains with high spatiotemporal resolution. Together, sterol substitution and AzoSM exchange offer critical insights into the role of cholesterol and sphingomyelin-dependent lipid domains in mediating membrane receptor signaling. Interestingly, protein tyrosine phosphatases have been reported to be excluded from lipid rafts during antigen-activated IgE signaling mediated by Fcε receptor I (FcεRI) (66). In addition, a loss of insulin receptor accessibility to phosphatases when lipid raft levels are increased regulates its kinase activity (37). Consequently, exclusion of phosphatases from raft domains provides an amenable environment for efficient kinase activity and IgE signaling. Since FcγRIII and FcεRI both require FcRγ phosphorylation to initiate the signaling cascade (67), raft disruption in our studies may have resulted in aberrant phosphatase activity on FcRγ to attenuate IgG signaling.

Together these data demonstrate that lipid raft formation, high membrane order, and domain structure are critical for FcγRIII function whereby disruption of lipid rafts interferes with the efficiency of FcγRIII signaling. Through this work, we provide direct evidence that cholesterol and sphingomyelin play a critical role in coordinating a phagocytic response to *C. neoformans*.

## Experimental procedures

### Cells, media, and reagents for lipid alteration

*C. neoformans* var. *grubii* serotype A strain H99 was used in this study. Fungal cells were grown in yeast nitrogenous base (YNB) medium containing 2% glucose, pH 7.2, for 16 to 18 hours at 30°C with shaking before experiments. Alveolar macrophage cell line MH-S (ATCC CRL-2019) was cultured in complete RPMI-1640 (supplemented with 10% FBS and 1% penicillin/streptomycin) at 37°C + 5% CO_2_. Peritoneal macrophage cell line J774.1 (ATCC TIB-67) was cultured in complete high glucose DMEM (supplemented with 10% FBS and 1% penicillin/streptomycin). Macrophages were seeded at the following densities for experiments: 5 x 10^4^ cells in 200 μl per well for 96-well plates (phagocytosis assay), 1 x 10^6^ (lipid analysis) or 1.5 x 10^6^ (flow cytometry) in 5 ml per well for 6-well plates, 1 x 10^6^ (microscopy) in 5 ml per 35 mm glass bottom dish, and 8.6 x 10^6^ (immunoprecipitation/Western blot) or 1.5 x 10^7^ (FRET) in 15 ml for 100 mm dishes. Cells were allowed to adhere at 37°C + 5% CO_2_ overnight prior to commencing experiments. AzoSM was synthesized as previously described (55). Sphingomyelinase from *Staphylococcus aureus* (Sigma), methyl-beta-cyclodextrin (MβCD) (Sigma), cholesterol (Sigma), 7-dehydrocholesterol (Sigma), and coprostanol (Cayman Chemicals) used for lipid altering treatments were purchased.

### Preparation of sterol-loaded methyl-beta-cyclodextrin (MβCD)

Sterol-loaded MβCD solutions were prepared as previously described (32,33). Briefly, sterols dissolved in chloroform were dried under nitrogen gas (N-EVAP, Organomation) followed by high vacuum for at least 1 hour (Savant SpeedVac, Thermo Fisher Scientific). They were then hydrated with 2.5 mM MβCD prepared in serum-free medium. Sterol-loaded MβCD solutions were then sonicated (Bransonic Ultrasonic Baths, Branson) for 5 minutes until no visible particles remained. Next, the sterol-loaded MβCD solutions were incubated at 55°C for 30 minutes then at 37°C overnight with shaking. Solutions were filtered through a 0.22 μm polyethersulfone syringe filter (Millex-GP, Millipore Sigma) to remove undissolved sterol. Unless otherwise noted, the final solutions, assuming undissolved sterol was negligible, contained 0.2 mM cholesterol, 7-dehydrocholesterol, or coprostanol mixed with 2.5 mM MβCD.

### Cholesterol depletion/sterol repletion using MβCD

Cholesterol was depleted from macrophages using MβCD as previously described (38). Briefly, macrophages were seeded as described above and allowed to adhere overnight at 37°C + 5% CO_2_. Macrophage monolayers were washed twice with 1x Dulbecco’s phosphate buffered saline (DPBS). 10 mM or 30 mM MβCD solutions were prepared in serum-free media (DMEM for J774.1 cells and RPMI-1640 for MH-S cells) and added to macrophage monolayers as follows: 200 μl for 96-well plates, 2 ml for 6-well plates or 35 mm dishes, and 5 ml for 100 mm dishes. Cells were incubated in MβCD solution for 30 minutes at 37°C + 5% CO_2_ with shaking. After removing the MβCD solution, cells were washed three times with 1x DPBS before processing for analysis.

For sterol repletion, cholesterol was first depleted from macrophages using 10 mM MβCD in serum-free medium for 30 minutes 37°C + 5% CO_2_ with gentle shaking. The macrophage monolayers were then washed three times with 1x DPBS. Sterol-loaded MβCD solutions were added to the cells as follows: 200 μl for 96-well plates, 1 ml for 6-well plates or 35 mm dishes, and 2 ml for 100 mm dishes. Cells were incubated at 37°C + 5% CO_2_ for 2 hours with gentle shaking. Macrophage monolayers were then washed four times with 1x DPBS before processing for analysis.

### Sphingomyelin depletion using bacterial sphingomyelinase

To reduce sphingomyelin (SM) in the outer leaflet of the plasma membrane, macrophages were treated with bacterial sphingomyelinase (bSMase) to catalyze the transformation of sphingomyelin into ceramide and phosphorylcholine as previously described (40). Briefly, macrophages were seeded as described above and allowed to adhere overnight at 37°C + 5% CO_2_. Macrophage monolayers were washed twice with 1x DPBS. 50, 100, 250, or 500 mU/ml bSMase solutions were prepared in serum-free media and added to macrophage monolayers as follows: 200 μl for 96-well plates, 2 ml for 6-well plates or 35 mm dishes, and 5 ml for 100 mm dishes. Cells were incubated in bSMase solution for 20 minutes at 37°C + 5% CO_2_ with shaking. Macrophage monolayers were then washed three times with 1x DPBS before processing for analysis.

### Preparation of AzoSM loaded methyl-alpha-cyclodextrin (MαCD)

AzoSM was loaded into MαCD for sphingomyelin exchange as previously described (32,36). Briefly, AzoSM dissolved in chloroform was dried under nitrogen gas (N-EVAP, ORganomation). The lipid was then resuspended in pre-warmed serum-free medium, vortexed vigorously, and incubated at 70°C for 5 minutes on a heat block. Next, the appropriate amount of MαCD was added to the solution to yield 40 mM MαCD. To load AzoSM into MαCD, the solution was then incubated for 30-45 minutes at 37°C + 5% CO_2_ with constant shaking until no visible particles remained. The final solution, assuming undissolved AzoSM was negligible, contained 1.5 mM AzoSM mixed with 40 mM MαCD.

### AzoSM exchange using MαCD and photoisomerization

Endogenous sphingomyelin was exchanged for AzoSM using MαCD as previously described (36,38). Briefly, macrophages were seeded as described above and allowed to adhere overnight at 37°C + 5% CO_2_. Macrophage monolayers were washed twice with 1x Dulbecco’s phosphate buffered saline (DPBS). AzoSM-loaded MαCD solution was then added to the cells as follows: 200 μl for 96-well plates, 1 ml for 6-well plates, and 2 ml for 100 mm dishes. Cells were incubated at 37°C + 5% CO_2_ for 1 hour with gentle shaking. Macrophage monolayers were then washed four times with 1x DPBS and incubated in serum-free media. To isomerize AzoSM from the dark-adapted *trans* conformation to the *cis* conformation, cells were treated with UV light (λ= 365 nm) for 20 seconds. To revert the isomerization (from *cis* conformation back to *trans* conformation), cells were subsequently treated with blue light (λ= 460 nm) for 40 seconds. Cells treated with UV light only subsequently remained in the dark to minimize isomerization back to *trans*.

### In vitro phagocytosis assay

5 x 10^4^ macrophages were seeded per well in 96-well plates. Cells were allowed to adhere overnight at 37°C + 5% CO_2_. *C. neoformans* H99 cells were grown overnight in YNB at 30°C with shaking. Fungal cells were then washed twice with 1x phosphate buffered saline (PBS). For antibody-mediated phagocytosis, *C. neoformans* H99 cells were opsonized for 30 minutes at 37°C + 5% CO_2_ with shaking using 1 μg of anti-glucuronoxylomannan (GXM) antibody (39) per 10^5^ *C. neoformans* cells in 100 μl of complete media (10% FBS + 1% penicillin/streptomycin). Antibody-opsonized *C. neoformans* cells were added to washed macrophage monolayers at a 1:1 ratio in 200 μl and allowed to interact for 2 hours at 37°C + 5% CO_2_ unless noted otherwise. Macrophage monolayers were then washed three times with 1x DPBS to remove extracellular *C. neoformans*, fixed with ice-cold methanol for 15 minutes at room temperature (RT), and stained for 5 minutes at RT with Giemsa (Sigma) diluted 1:1 in deionized water. Stained cells were then washed three times with sterile deionized water and allowed to dry overnight at room temperature. For complement-mediated phagocytosis assays, *C. neoformans* H99 cells were opsonized for 1 hour at 37°C + 5% CO_2_ with shaking using 100 μl of complement-based media (20% mouse CD1 complement serum (Innovative Research) in serum-free medium). Complement-opsonized *C*. neoformans cells were added to washed macrophage monolayers at a 1:1 ratio with 10 units (1 unit = 0.1 ng/mL) of recombinant murine IFNγ (Sigma) and 0.3 μg/ml of lipopolysaccharide (Sigma) in 200 μl and allowed to interact for 3 hours at 37°C + 5% CO_2_. Macrophage monolayers were then washed three times with 1x DPBS to remove extracellular *C. neoformans*, fixed with ice-cold methanol for 15 minutes at RT, and stained for 5 minutes at RT with Giemsa (Sigma) diluted 1:1 in deionized water. Stained cells were then washed three times with sterile deionized water and allowed to dry overnight at room temperature. Phagocytic index was calculated by microscopic observation. Micrographs were taken of at least 4 independent fields of view per well. At least 200 macrophage cells were counted per well. Ingested *C. neoformans* cells within macrophages were counted. Ingested *C. neoformans* cells were distinguished from attached cells by the stained macrophage cell membrane surrounding the cryptococcal capsule. Phagocytic index is defined as the percentage of yeast cells ingested per macrophages counted per well (9).

### Lipid analysis

Sphingomyelin and ceramide species were analyzed as previously described (75). 1 x 10^6^ macrophages were seeded per well in 6-well plates. Cells were allowed to adhere overnight at 37°C + 5% CO_2_. Macrophage monolayers were washed twice with 1x DPBS and treated for sphingomyelin depletion as described above. Following treatment, cells were washed three times with 1x DPBS and subsequently scraped in 1 ml ice-cold 1x DPBS. 10 μl of cell suspension was set aside for protein quantification using the Bradford method. Remaining cells were pelleted at 1,000 x g for 5 minutes at 4°C. Lipids were extracted from each cell pellet in 2 ml isopropanol:water:ethyl acetate (30:10:60 by volume). Cell extracts were analyzed by reverse phase high pressure liquid chromatography coupled to electrospray ionization and subsequent separation by mass spectrometry. Analysis of ceramides and sphingomyelins was performed on a TSQ Quantum Ultra mass spectrometer (Thermo Fisher Scientific), operating in a multiple reaction-monitoring positive ionization mode. Sphingolipid levels were normalized to the protein concentration estimated in the original cell extract. All experiments were performed in triplicates.

For analysis of sterols, 1 x 10^6^ macrophages were seeded per well in 6-well plates. Cells were allowed to adhere overnight at 37°C + 5% CO_2_. Macrophage monolayers were rinsed twice with 1x DPBS and treated for cholesterol depletion or sterol substitution as described above. Following treatment, cells were washed three times with 1x DPBS and subsequently scraped in 1 ml ice-cold 1x DPBS. 10 μl of cell suspension was set aside for protein quantification using the Bradford method. Remaining cells were pelleted at 1,000 x g for 5 minutes at 4°C. Sterol derivatization, detection and analysis were performed by gas chromatography–mass spectrometry (GC-MS) as previously described (33,76). Briefly, lipids were first extracted in 2 ml isopropanol:water:ethyl acetate (30:10:60 by volume), then subjected to mild alkaline base hydrolysis (0.6 M KOH in methanol) to remove the glycerol backbone containing lipid contaminants (75,77). The extracted and base hydrolyzed lipid samples were derivatized using 100 μl BSTFA/TMCS (N,O-bis (trimethylsilyl) trifluoroacetamide/TMCS (trimethylchlorosilane)) reagent at 85°C for 90 min. Next, 50 μl n-hexane was added to the derivatized sample and vortexed. The samples were then analyzed using 30 meter (0.25 μm) DB5-MS column on Agilent 7890 GC-MS (Agilent Technologies). The initial column temperature of 10°C was held for 0.5 min, ramped up at 35°C/min to 240°C, then at 3°C/min to 260°C, and then at 1.5°C/min to 305°C with hold of 2 min. All EI-mass spectra were recorded at 70 eV with ion source temperature of 230°C. Front inlet temperature was kept at 290°C and MSD transfer line temperature was kept at 280°C. The retention time and mass spectral patterns of the appropriate sterol standards were used as a reference. Ergosterol (25 μg), a non-mammalian sterol, was added as an internal standard for these analyses prior to lipid extraction. Sterol species contents in each sample were quantified using standard calibration curves and normalized to protein concentration estimated in the original cell extract. All experiments were performed in triplicates.

For AzoSM analysis, 1 x 10^6^ macrophages were seeded per well in 6-well plates. Cells were allowed to adhere overnight at 37°C + 5% CO_2_. Macrophage monolayers were rinsed twice with 1x DPBS and treated for AzoSM exchange as described above. Following treatment, cells were washed four times with 1x DPBS and subsequently scraped in 1 ml ice-cold 1x DPBS. 10 μl of cell suspension was set aside for protein quantification using the Bradford method. Remaining cells were evenly split into two tubes and pelleted at 1,000 x g for 5 minutes at 4°C. One tube was set aside for sphingomyelin analysis as described above. For AzoSM analysis, cellular lipids were extracted from the second tube for each sample in 1.5 ml Mandala extraction buffer (78), followed by Bligh and Dyer extraction (79), and base hydrolysis (77). Cell extracts were each resuspended in 50 μl methanol and 10 μl applied to high-performance liquid chromatograph (HPLC) (Agilent Technologies). AzoSM was detected at 365 nm on a C-8 column with a flow rate of 0.5 ml/min in methanol/water 90:10 ratio buffered with 1mM ammonium formate and 0.2% formic acid (Figure S8). Total AzoSM was calculated for each sample, then normalized to the protein concentration estimated in the original cell extract. All experiments were performed in triplicates.

### Förster resonance energy transfer (FRET)

Biological membrane nanodomain stability was assessed by measuring the temperature dependence of FRET using giant plasma membrane vesicles (GPMVs) as previously described (36). 1.5 x 10^7^ macrophages were seeded in 100 mm dishes. Cells were allowed to adhere overnight at 37°C + 5% CO_2_. Macrophage monolayers were rinsed twice with 1x DPBS and treated for cholesterol depletion or sterol substitution as described above. Following treatment, GPMVs were prepared using as previously described by Sezgin et al. (80) and Levental and Levental (80,81). Briefly, paraformaldehyde (PFA) and dithiothreitol (DTT) were freshly added to the GPMV buffer (10 mM HEPES, 150 mM NaCl, 2 mM CaCl_2_, pH 7.4) at a final concentration of 24 mM PFA and 2 mM DTT when desired. Following cholesterol depletion or sterol substitution, macrophage monolayers were washed twice with 1x DPBS then twice with GPMV buffer lacking PFA and DTT. Next, 2.5 ml of GPMV buffer with PFA and DTT was added and incubated at 37°C + 5% CO_2_ with gentle shaking. The GPMV-containing buffer solution was gently harvested by pipetting. To remove intact cells, the solution was centrifuged at 100 x g for 5 minutes at RT. This GPMV-containing buffer solution was used for the FRET measurements. FRET measurements were made with diphenylhexatriene (DPH) (Sigma) as the FRET donor and Octadecyl Rhodamine B (ODRB) (Invitrogen) as the FRET acceptor. The “F sample” with FRET acceptor was prepared by adding 3.6 μl from 1.4 mM ODRB dissolved in ethanol to 1 ml of GPMVs, vortexed, and incubated at 37°C for 1 hour in the dark. The “F_o_ sample” lacking FRET acceptor was also incubated at 37°C for 1 hour with 3.6 μl ethanol. Before fluorescence measurements, DPH (1.2 μl from a 5 μM stock solution in ethanol) was added to both the “F sample” and “F_o_ sample” and incubated at room temperature for 5 min in the dark. Background fluorescence before adding DPH was negligible (<1% of samples containing DPH). To initiate measurements, samples were placed in a cuvette and transferred to a temperature-controlled sample spectrofluorometer (Horiba PTI Quantamaster) sample holder and cooled to 16°C. DPH fluorescence (excitation λ=358 nm, emission λ=430 nm) was measured. Temperature was increased at 4°C intervals every 5 minutes up to 64°C. The ratio of DPH fluorescence intensity in the presence of ODRB to that in its absence (F/F_0_) was then calculated. All F/F_0_ values were normalized to the F/F_0_ value measured at 64°C. T_end_, the temperature above which segregation of lipids into ordered and disordered domains was fully lost (i.e., complete “melting” of ordered domains) was estimated by finding the minimum value of a polynomial fit applied to the F/F_0_ as previously described (36). Presence of detectable nanodomains between 16°C and T_end_ was estimated by calculating the area between the polynomial fit and a horizontal line defined by T_end_ (i.e. y = y value at x = T_end_ for polynomial fit). All FRET experiments were performed three times.

### Flow cytometry

Flow cytometry was used to assess the presence of Fcγ receptors (CD16/CD32) and complement receptor (CD11b) on the cell surface following lipid-altering treatments. 1 x 10^6^ macrophages were seeded per well in 6-well plates. Cells were allowed to adhere overnight at 37°C + 5% CO_2_. Macrophage monolayers were rinsed twice with 1x DPBS and treated for cholesterol or sphingomyelin depletion as described above. Following treatment, cells were washed three times with 1x DPBS and subsequently scraped in 1 ml icecold 1x DPBS. Cell concentrations were determined using a hemacytometer. 1 x 10^6^ cells were transferred into round-bottom test tubes (Falcon) per experimental sample and compensation control (i.e. unstained, single stained) and washed twice with 1x PBS + 2% FBS. Cells were labeled with the following antibodies directly conjugated to the following fluorophores: CD45-PE/Cy7 (clone 30-F11, BioLegend), CD11b-BV510 (clone M1/70, BioLegend), and CD16/CD32-APC (clone 93, Invitrogen). Cells were also stained for viability using Alexa Fluor 700 carboxylic acid, succinimidyl ester (Thermo Fisher Scientific). Additional details for the antibodies/markers are available in Table S2. Cells were first incubated with CD16/CD32-APC at 4°C for 30 minutes in the dark, then incubated with CD11b-BV510, CD45-PE/Cy7, and the viability stain at 4°C for 30 minutes in the dark. All labeling was performed with shaking. Cells were then washed twice with 1x PBS + 2% FBS and fixed with Fluorofix Buffer (BioLegend) at RT for 30 minutes in the dark with shaking. Cells were washed once and resuspended in 400 μl 1x PBS + 2% FBS for analysis. Labeled cell suspensions were analyzed using the CytoFLEX Flow Cytometer (Beckman-Coulter) and FlowJo Software Version 10.6.2 to measure the median fluorescence intensity of CD16/CD32-APC, CD11b-BV510, and CD45-PE/Cy7 for 50,000 live CD45^+^ cells. All experiments were performed in triplicates.

### Fcγ receptor stimulation using IgG immune complexes

To assess activation of the Fcγ receptor signaling cascade, macrophages were stimulated with BSA-IgG immune complexes (IgGIC) (59,82). Briefly, IgGIC was prepared by incubating 500 μg of anti-BSA IgG (MP Biomedicals) per 100 μg of BSA (fatty acid-free, protease-free, Sigma) in 100 μl 1x PBS at RT for 30 minutes with shaking. IgGIC was diluted with complete media to a final concentration of 100 μg/ml. 8.6 x 10^6^ macrophages were seeded in 100 mm dishes. Cells were allowed to adhere overnight at 37°C + 5% CO_2_. Macrophage monolayers were rinsed twice with 1x DPBS and treated for cholesterol depletion, sterol substitution, or sphingomyelin depletion as described above. Following treatment, cells were washed three times with 1x DPBS and incubated with 2.5 ml IgGIC at 37°C + 5% CO_2_ with gentle shaking. Cell were then washed four times with 1x DPBS before processing for immunoprecipitation and immunoblotting.

### Immunoprecipitation and immunoblotting

Following Fcγ receptor stimulation using IgGIC, Fc receptor γ-subunit (FcR γ subunit) was immunoprecipitated and assessed for phosphorylation of tyrosine residues (59). Cells were lysed directly on the 100 mm dishes using icecold radio-immunoprecipitation assay (RIPA) buffer (Sigma) supplemented with 0.1% Triton X-100 (Sigma) and Halt Protease and Phosphatase Inhibitor Cocktail (Sodium Fluoride, Sodium Orthovanadate, β-glycerophosphate, Sodium Pyrophosphate, Aprotinin, Bestatin, E64, Leupeptin) (Thermo Fisher Scientific). Samples were incubated at 4°C for 45 minutes with shaking. Samples were then centrifuged at 4°C, 10,000 rpm for 5 minutes and supernatants were transferred to pre-chilled microtubes. Protein concentration of each sample was determined by the Bradford assay. 200 μg each lysate was pre-cleared with 20 μl of Protein A agarose beads (3 mg/ml, Roche) in 200 μl for 1 hour at 4°C on a Labquake Rotator (Barnstead Thermolyne). Pre-cleared lysate was diluted to 500 μl with additional lysis buffer and incubated overnight with 40 μl Protein A agarose beads and 4 μg anti-FcRγ-subunit IgG (Millipore Sigma) at 4°C on a Labquake Rotator. Beads were then centrifuged at 4°C, 10,000 rpm for 5 minutes and rinsed five times with lysis buffer. Beads were then mixed with 40 μl of Laemmli Sample Buffer (BioRad) containing 5%β-mercaptoethanol (Sigma) and boiled for 5 minutes at 95°C. All samples were centrifuged briefly at 14,000 rpm before being loaded onto two SDS-polyacrylamide gel (4%-15% gradient gel, BioRad). Precision Plus Protein Kaleidoscope pre-stained protein ladder (BioRad) was loaded along with the samples. After separation by electrophoresis at 100 V for 60 minutes in Tris-Glycine/SDS buffer, proteins were transferred to Immobilon-PSQ PVDF membrane (0.2 μm pore size, Millipore Sigma) by electrophoresis at 100 V for 30 minutes in Tris-Glycine/Methanol buffer. Nonspecific antibody binding was blocked using 10% non-fat dried milk (NFDM) or 10% BSA in 1x Tris-buffered saline + 0.05% Tween 20 (1x TBST).

After blocking with NFDM/1x TBST, FcR γ subunit was probed with the anti-FcR γ subunit IgG (Millipore Sigma) overnight at 4°C with gentle shaking. After blocking with BSA/1x TBST, phosphor-tyrosine was probed with the anti-phospho-tyrosine MultiMab mix (Cell Signaling) 4°C with gentle shaking. After incubation with primary antibody, membranes were washed five times with 1x TBST. Both membranes were subsequently incubated for 1 hour at RT with the Clean-Blot IP Detection Reagent (HRP, Thermo Scientific). After washing the membrane three times with 1x TBST, the secondary antibody was detected using SuperSignal West Femto Maximum Sensitivity Substrate (Thermo Scientific). Bands were visualized using ImageQuant LAS 500 (Cytiva). Bands were quantified using ImageJ. Phosphotyrosine bands were normalized by FcR γ subunit bands. All experiments were performed three times. Additional details for the antibodies are available in Table S2.

### Fluorescent lipid labeling

EGFP-conjugated nakanori (EGFP-nakanori), a protein that labels cell surface domains in a sphingomyelin and cholesterol-dependent manner (46), was used to label macrophages. Plasmid containing EGFP-nakanori was transformed into *Escherichia coli* strain BL21. Expression was induced in LB media containing Isopropyl β-D-1-thiogalactopyranoside. The EGFP-nakanori protein was subsequently extracted and purified using HisTrap columns (Cytiva) and concentrated using an Amicon centrifugal unit.

Macrophages were subjected to either 10 mM MβCD or 250 mU/ml bSMase treatment as described above. Live cells were subsequently coincubated with 50 μg EGFP-nakanori for 1 hour, fixed with 4% formaldehyde, and imaged using the Axio Observer D1 Phase Contrast Fluorescence Microscope (Zeiss). Micrographs were captured using the AxioCam MR R3 camera (Zeiss).

### Statistical analysis

For all statistical analysis, GraphPad Prism Version 8 software was used. Data are represented as mean ± standard error of the mean (SEM). α-level (type 1 error) was set at 0.05. Differences were considered significant when the probability of type 1 error was less than 5% (P < 0.05). To compare groups, one-way analysis of variance (ANOVA) with Tukey’s multiple comparisons post hoc test was used. All P values from multiple comparisons post hoc tests were corrected for multiplicity using the Bonferroni adjustments.

## Data availability

All data are contained within the manuscript.

## Supporting information

This article contains supporting information.

## Acknowledgements

We would like to acknowledge Stony Brook Cancer Center (Biological Mass Spectrometry Shared Resource) for expert assistance with sphingolipid analysis and the technical support provided by the Research Flow Cytometry Laboratory at Stony Brook Medicine. We especially thank Izolda Mileva and Todd Rueb. We acknowledge Arturo Casadevall from Johns Hopkins University (Baltimore, Maryland, USA) for providing the anti-GXM antibody (clone 18B7) used for opsonization of *C. neoformans* and Yasushi Sako from RIKEN BioResource Research Center (Kyoto, Japan) for providing the EGFP-nakanori plasmid used for cell labeling. We would also like to acknowledge the contributions of Chiara Luberto for her insightful discussion and set up of the initial experiments.

## Author contributions

AMB and JKY conceptualized and performed the experiments, collected the data, performed statistical analyses, and wrote the manuscript. GL helped with FRET experiments and AzoSM exchange protocol. JHK helped with the sterol substitution protocol. AS helped with GC-MS. JM and DT synthesized AzoSM. NPS helped with HPLC. TGN helped with Nakanori staining for microscopy. AF, EL, and MDP conceptualized the experiments and discussed the data. All authors edited the manuscript. The order of co-first authors was determined alphabetically by last name.

## Funding and additional information

This work was supported by NIH grants AI136934, AI116420, and AI125770 to MDP and GM122493 to EL, the Merit Review Grant I01BX002924 from the Veterans Affairs Program to MDP, Department of Defense grant PR190642. MDP is a recipient of the Research Career Scientist (RCS) Award (IK6 BX005386) and a Burroughs Welcome Investigator in Infectious Diseases at the Veterans Administration Medical Center in Northport, NY. The content is solely the responsibility of the authors and does not necessarily represent the official views of the National Institutes of Health.

## Abbreviations

(bSMase): bacterial sphingomyelinase
(IgGICs): BSA-IgG immune complexes
(FcRγ): Fc receptor γ-subunit
(FcγR): Fcγ receptor
(FcεRI): Fcε receptor I
(FRET): Förster resonance energy transfer
(GPMVs): giant plasma membrane vesicles
(GXM): glucuronoxylomannan
(ITAM): immunoreceptor tyrosine-based activation motif
(ITIM): immunoreceptor tyrosine-based inhibitory motif
(MβCD): methyl-beta-cyclodextrin

---

**Table S1.** Nanodomain characteristics for GPMVs after cholesterol depletion or sterol repletion

| *Treatment condition* | *T<sub>end</sub> (°C)* | *Relative nanodomain formation (area)* |
| --- | --- | --- |
| Untreated control | 50.6 ± 0.7 | 18.1 ± 1.1 |
| Depletion: 10 mM MβCD | 45.1 ± 0.8 | 9.6 ± 1.3 |
| Depletion: 30 mM MβCD | 39.8 ± 6.4 | 2.2 ± 1.1 |
| Repletion: 0.2 mM cholesterol/2.5 mM MβCD | 52.3 ± 1.6 | 28.0 ± 3.7 |
| Repletion: 0.2 mM 7-dehydrocholesterol/2.5 mM MβCD | 56.2 ± 1.7 | 42.2 ± 2.2 |
| Repletion: 0.2 mM coprostanol/2.5 mM MβCD | 41.3 ± 0.9 | 2.7 ± 0.5 |
| 250 mU/ml bSMase treatment | 54.9 ± 1.7 | 17.6 ± 1.9 |
| 500 mU/ml bSMase treatment | 52.2 ± 2.0 | 14.6 ± 3.5 |

Mean and standard error of the mean (SEM) values from three experiments are shown, except for untreated controls (n=9) and depletion with 10 mM MβCD (n=6).

**Table S2.** Antibodies/markers used in this study

| Catalog number | Manufacturer | Name | Final dilution | Experiment |
| --- | --- | --- | --- | --- |
| 103114 | BioLegend | CD45-PE/Cy7 | 1:800 | Flow cytometry |
| 101263 | BioLegend | CD11b-BV510 | 1:500 | Flow cytometry |
| 17-0161-82 | Invitrogen | CD16/32-APC | 1:800 | Flow cytometry |
| A20110 | Invitrogen | Alex Fluor 700, carboxylic acid, succinimidyl ester | 1:1200 | Flow cytometry |
| 06-727 | Sigma-Aldrich | anti-FcεRI, $\gamma$ subunit | 1:1000 | Western blot |
| 8954S | Cell Signaling Technology | Phospho-Tyrosine (P-Tyr-1000) MultiMab™ Rabbit mAb mix | 1:3000 | Western blot |
| 21230 | Thermo Scientific | Clean-Blot™ IP Detection Reagent (HRP) | 1:1000 | Western blot |

